# NRF2 inhibition of alveolar macrophage MHC II expression during *Mycobacterium tuberculosis* infection

**DOI:** 10.1101/2025.08.16.670319

**Authors:** Linh K. Pham, Maritza M. Cervantes, Pamelia N. Lim, Divya Dubey, Ankita Tufts, Tomoyo Shinkawa, Samuel M. Behar, Alissa C. Rothchild

## Abstract

During *Mycobacterium tuberculosis* (Mtb) infection, infected alveolar macrophages (AMs) initially up-regulate a NRF2 regulated cell-protective program, which is detrimental to host control and impedes AM activation, including MHC II expression. MHC II is critical for CD4^+^ T cell activation and host immunity during Mtb infection. We hypothesized that NRF2 regulates the MHC II pathway and AM antigen presentation to T cells. We found that NRF2 inhibits MHC II, but not MHC I, specifically in AMs, following Mtb infection *in vitro* and *in vivo*. NRF2 dampens *Ciita* and *H2-Ab1* gene expression in uninfected AMs, and MHC II inhibition by NRF2 is retained following innate stimuli and IFNγ exposure. NRF2 expression in Mtb-infected AMs impedes their ability to activate ESAT6-specific CD4^+^ T cells. Thus, although NRF2 expression enhances cell-protective functions, it has the unexpected consequence of limiting innate-adaptive crosstalk, which can impair CD4+ T cell activation and host immunity during Mtb infection.

**HIGHLIGHTS:**

- NRF2 inhibits MHC II on AMs, but not DCs, MDMs, or PMNs, during Mtb infection
- NRF2 inhibits *Ciita* and *H2-Ab1* and total MHC II protein in AMs, but not in BMDMs
- NRF2 blockade of AM MHC II is retained following PAMP and IFNγ stimulation
- NRF2 impedes the activation of antigen-specific CD4^+^ T cells by Ams

## INTRODUCTION

Tuberculosis (TB), a respiratory disease caused by the bacteria *Mycobacterium tuberculosis* (Mtb), remains a formidable global health issue. In 2023, Mtb was responsible for 1.25 million deaths and 10.8 million new cases^1^. Antibiotic therapy can be highly effective against TB, but existing treatments are lengthy, costly, and can be highly toxic. Ultimately, existing drug treatments fall short of controlling the TB epidemic. Additionally, incomplete drug treatment can lead to multidrug-resistant Mtb. Host-directed therapies, including vaccines, are needed to improve host responses to Mtb and protect against disease.

Alveolar macrophages (AMs) are the first cells in the lung to be infected with Mtb after aerosol transmission^2,3^. AMs are long-lived lung-resident myeloid cells, with a turnover rate of just 20% in 12 weeks^4^ or ∼40% in 1 year^5^. AMs serve critical homeostatic functions in the lung that include clearance of debris from the airways and surfactant recycling^6,7^. Mtb-infected AMs promptly initiate a cell-protective and antioxidant transcriptional program, regulated by the transcription factor nuclear factor erythroid 2– related factor 2 (NRF2)^3^. Up-regulation of the NRF2 transcriptional program within Mtb-infected cells requires the pulmonary microenvironmental^3^. Despite serving as the initial innate sentinels in the airway, AMs exhibit a diminished capacity to control bacterial growth when compared to recruited monocyte-derived macrophages (MDMs)^8^. However, AMs that lacked NRF2 expression exhibited better control of Mtb over the first 10 days of infection than WT AMs^3^. Interestingly, over the same time frame, AMs deficient in NRF2 expressed higher levels of the antigen presentation protein, MHC II^3^. MHC II is necessary for antigen-presenting cells to interact with CD4^+^ T cells and induce effector T cell responses^9,10^. In the context of respiratory Mtb infection, tissue-resident memory T cells are critical for control of Mtb^11^ and are known to colocalize with and receive signals from tissue-resident macrophages around the airways in human lungs^12^. We hypothesized that NRF2 regulates the ability of AMs to interact with CD4^+^ T cells through MHC II.

While NRF2 has well-known roles in the cellular defense against oxidative stress and inflammation^13^, its role in antigen presentation has not yet been explored. NRF2 regulates numerous genes involved in antioxidant and detoxification processes and is a target of cancer therapy^14–17^ and respiratory inflammation, such as COPD^18,19^. NRF2 activity is tightly regulated through interaction with the Kelch-like ECH-associated protein 1 (KEAP1), which targets NRF2 for proteasomal degradation under baseline conditions. Oxidative stress or electrophilic compounds can disrupt this intracellular interaction, allowing NRF2 to dissociate from KEAP1, translocate into the nucleus, and promote transcription of antioxidant or anti-inflammatory genes^20^. NRF2 has been shown to suppress pro-inflammatory cytokine transcription induced by LPS stimulation in bone marrow-derived macrophages (BMDMs)^21^ and block ferroptosis, a form of cell death dependent on lipid peroxidation and iron^22–25^. Despite these broad effects, NRF2 regulation of antigen presentation is unknown.

To investigate the importance of NRF2-mediated inhibition of MHC class II expression in AM, we used an *in vivo* murine aerosol infection model and *in vitro* murine macrophage models, including primary AMs and *ex vivo* murine AMs (mexAMs)^26^ derived from WT, NRF2 KO (NRF2^-/-^), or NRF2 myeloid-conditional knock-out strains as well as BMDMs, as a model of recruited macrophages. We show that both *in vivo* and *in vitro*, NRF2 inhibits expression of MHC II on AMs, but not dendritic cells (DCs) or other macrophages. In addition to blocking MHC II surface expression, NRF2 inhibits *Ciita* and *H2-Ab1* gene expression and total MHC II protein. We find that inhibition of MHC II expression by NRF2^-/-^ is retained after stimulation with innate Pathogen Associated Molecular Patterns (PAMPs) or IFNɣ. Lastly, we demonstrate that, in the absence of NRF2, AMs have an enhanced ability to activate ESAT6-specific C7 TCR transgenic CD4^+^ T cells in a macrophage-T cell co-culture system. Overall, our results demonstrate that NRF2 impedes the gene and protein expression of MHC II specifically in AMs, which ultimately blocks their ability to activate CD4^+^ T cells in an antigen-specific manner.

## RESULTS

### NRF2 limits MHC II expression on AMs during Mtb infection

To determine the effect of NRF2 on MHC II expression of Mtb-infected AM *in vivo*, we used two different myeloid NRF2 conditional knock-out (NRF2^CondKO^) strains Nrf2^fl/fl^LysM^cre^ (NRF2 deleted in AMs, MDMs and neutrophils (PMNs)) and Nrf2^fl/fl^CD11c^cre^(NRF2 deleted in AMs and DCs). We infected the NRF2^CondKO^ mice and their littermate control (Nrf2^fl/fl^) with fluorescent trackable Mtb strains (mCherry-H37Rv, msfYFP-H37Rv) and collected mouse lungs at day 12 and 14 days post-infection (dpi). The median fluorescence intensity (MFI) of MHC II levels on bystander and Mtb-infected myeloid cell populations were measured by flow cytometry (**Fig 1A**, see gating scheme, **Fig S1A**). At 14 dpi, Mtb-infected AMs from Nrf2^fl/fl^LysM^cre^ and Nrf2^fl/fl^CD11c^cre^ mice expressed significantly higher surface levels of MHC II compared to their littermate controls **(Fig 1B, C)**. Bystander AMs also showed higher expression of MHC II (**Fig 1B, C**). This result was consistent with our previous findings, which showed that Mtb-infected AMs from NRF2^CondKO^ mice expressed higher levels of MHC II at 10 dpi than AMs from littermate controls^3^

**Figure 1:**
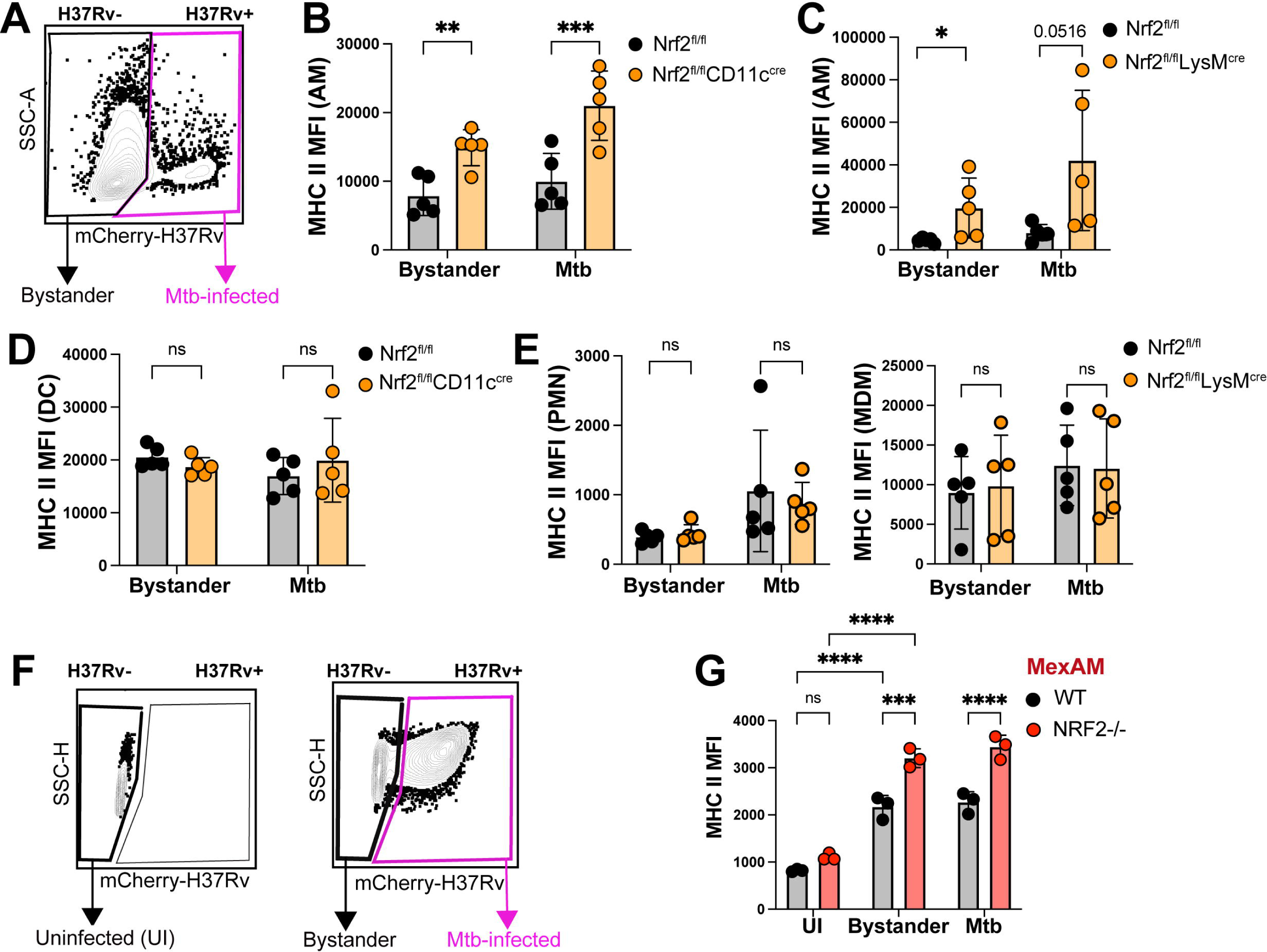
NRF2 limits MHC II expression on AMs during Mtb infection. **A)** Gating scheme for bystander and Mtb-infected AMs. **B, C)** MHC II MFI for Mtb-infected AMs and bystander AMs 14 dpi with mCherry-H37Rv from Nrf2^fl/fl^ CD11c^cre^ and Nrf2^fl/fl^ mice **(B),** or from Nrf2^fl/fl^ LysM^cre^ and Nrf2^fl/fl^ mice **(C). D)** MHC II MFI for Mtb-infected DCs and bystander DCs 14 dpi from Nrf2^fl/fl^ CD11c^cre^ and Nrf2^fl/fl^ mice. **E)** MHC II MFI for Mtb-infected PMNs and MDMs and bystander PMNs, MDMs 14 dpi in Nrf2^fl/fl^ LysM^cre^ and Nrf2^fl/fl^ mice. **F)** Gating scheme for uninfected, bystander, and Mtb-infected mexAMs 24 hours after mCherry-H37Rv infection (MOI 2.4:1). **G)** MHC II MFI for WT and NRF2^-/-^ mexAMs. **B-E)** Representative of 2 independent experiments. **G)** Representative of 3+ independent experiments. **B-E,G)** Two-way ANOVA with Tukey post-test. *p<0.05, **p< 0.01, ***p< 0.001, ****p< 0.0001.

We found that Mtb-infected and bystander DCs from Nrf2^fl/fl^CD11c^cre^ **(Fig 1D),** PMNs from Nrf2^fl/fl^LysM^cre^, and MDMs from Nrf2^fl/fl^LysM^cre^ **(Fig 1E)** did not show any difference in MHC II levels compared to myeloid cells from littermate controls. Interestingly, these results suggest that the impact of NRF2 on MHC II surface expression is specific for AMs and does not affect other pulmonary myeloid cells during the first 14 days of infection.

To determine whether we could replicate this phenotype *in vitro*, we infected mexAMs derived from WT and NRF2^-/-^ mice with mCherry-H37Rv for 24 hours and then measured MHC II **(Fig 1F**, see gating scheme **Fig S1B)**. Both Mtb-infected and bystander NRF2^-/-^ mexAMs had significantly higher expression of MHC II compared to WT mexAMs **(Fig 1G**). We repeated the infection with WT and NRF2^-/-^ BMDMs and both bystander and Mtb-infected NRF2^-/-^ BMDMs expressed similar levels of MHC II as WT BMDMs (see gating scheme, **Fig S1C)**. To test whether MHC II levels would be affected by exogenous activation of NRF2, we used a NRF2 agonist, L-sulforaphane (L-sulf). L-sulf binds to the cysteine residue of KEAP1, a NRF2 inhibitor, leading to KEAP1 dissociation from NRF2 and translocation of NRF2 into the nucleus^27^. L-sulf stimulation reduced MHC II expression in WT, but not NRF2^-/-^ BMDMs in both Mtb-infected and bystander populations (**Fig S1D**). Thus, in the context of exogenous NRF2 activation, NRF2^-/-^ BMDMs expressed significantly higher expression of MHC II than WT BMDMs (**Fig S1D**). Overall, our data demonstrate that NRF2 limits MHC II expression in AMs during *in vivo* and *in vitro* Mtb infection

### NRF2 impedes MHC II gene expression in AMs, but not BMDMs, at baseline

With the observation that NRF2 interferes with AM MHC II surface expression, we next investigated how NRF2 regulates MHC II expression. We used an *in vitro* culture model and the NRF2 agonist L-sulf to measure NRF2 activation in BMDMs and AMs. We observed that LPS stimulation was sufficient to induce NRF2 protein accumulation in the nuclei of AMs and addition of L-sulf did not further increase NRF2 staining (**Fig S2A**). In contrast, accumulation of NRF2 was only detected in LPS-stimulated BMDMs following the addition of L-sulf (**Fig S2B**). The levels of the NRF2 target, *Nqo1*, showed that *Nqo1* levels were high in AMs at baseline and not affected by L-sulf treatment, which validated the microscopy results (**Fig 2A**). In contrast, *Nqo1* expression was increased in BMDMs after the addition of L-sulf (**Fig 2B**). These results show that AMs have high baseline NRF2 activity, a characteristic that is not shared with BMDMs.

**Figure 2:**
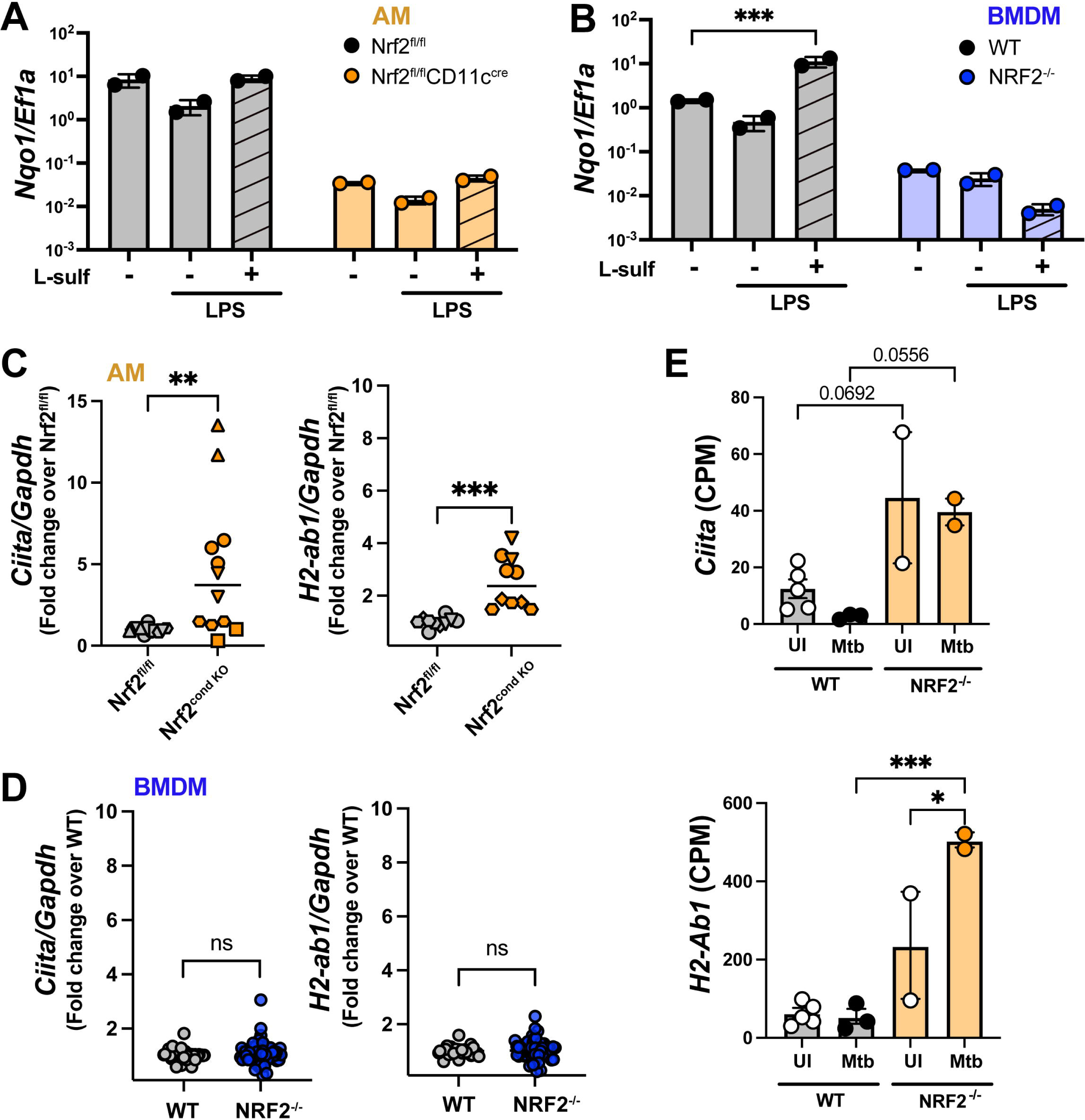
NRF2 impedes MHC II gene expression in AMs, but not BMDMs, at baseline. **A, B)** *Nqo1* expression after LPS stimulation with or without L-sulf, in Nrf2^fl/fl^ and Nrf2^condKO^ AMs (normalized to *Ef1a)* **(A)**, and in WT and NRF2^-/-^ BMDMs **(B)**. **C, D)** Baseline levels of *Ciita* and *H2-Ab1* expression (normalized to WT) in Nrf2^condKO^ and Nrf2^fl/fl^ AMs **(C),** and WT and NRF2^-/-^ BMDMs **(D)**. **E)** *Ciita* (left) and *H2-Ab1* (right) expression in naïve and Mtb-infected WT and NRF2^-/-^ AMs 24 hours post-infection, adapted from Rothchild et al, 2019^3^. **A-B, E)** Representative of 3 independent experiments, or **C, D)** Compiled from 4+ independent experiments per condition. **A,B)** Two-way ANOVA with Tukey post-test. **C, D)** Unpaired Student’s t-test. **E)** One-way ANOVA. *p <0.05, **p< 0.01, ***p< 0.001, ****p< 0.0001.

We examined regulation of the MHC II regulator, Class II transactivator (*Ciita)* ^9,28,29^, and *H2-Ab1,* one of many genes in the murine MHC II locus, by NRF2. We compared gene expression in AMs from Nrf2^CondKO^ mice (Nrf2^fl/fl^CD11c^cre^ and Nrf2^fl/fl^LysM^cre^) to Nrf2^fl/fl^ littermate controls. *Ciita* and *H2-Ab1* were significantly upregulated in Nrf2^CondKO^ AMs compared to Nrf2^fl/fl^ AMs (**Fig 2C**). Similarly, *H2-Ab1* expression was significantly higher in NRF2^-/-^ mexAMs compared to WT mexAMs, and a similar trend was observed for *Ciita* (**Fig S2C, D**). In contrast, *Ciita* and *H2-Ab1* levels were similar in WT and NRF2^-/-^ BMDMs (**Fig 2D**). Based on these *in vitro* results, we re-examined published transcriptional data from naive and Mtb-infected AMs sorted *ex vivo* from the lungs of WT and NRF2^-/-^ mice 24 hours following infection with high-dose mEmerald-H37Rv^3^. *Ciita* and *H2-Ab1* levels were elevated in both naive and Mtb-infected AMs from NRF2^-/-^ mice compared to AMs from WT mice (**Fig 2E**). Our findings reveal a distinctive regulatory role for NRF2 in suppressing baseline expression of MHC II genes in AMs. Consistent with the difference in baseline NRF2 activation, regulation of MHC II gene expression by NRF2 is present in AMs, but not other macrophage subsets, highlighting a potentially unique aspect of MHC II regulation within the alveolar microenvironment.

### NRF2 inhibits MHC II protein levels in AMs, but not BMDMs

Given that NRF2^-/-^ AMs expressed significantly higher levels of *Ciita* and *H2-Ab1* than WT AMs, we measured NRF2-regulated differences in MHC II internal and surface protein expression in AMs and BMDMs by flow cytometry (**Fig S3A, B, C, D**). Surface MHC II of uninfected Nrf2^fl/fl^LysM^cre^ AMs was significantly higher than Nrf2^fl/fl^ AMs **(Fig 3A)**. Compiling data over multiple experiments, we found that MHC II surface expression in both AMs and mexAMs lacking NRF2 expression was significantly higher (1.61-fold, 1.76-fold, respectively) than their respective WT controls (**Fig 3B**). In contrast, NRF2^-/-^ BMDMs showed only a moderate increase in MHC II surface expression (1.08-fold) compared to WT BMDMs (**Fig 3B**). In addition to transcriptional regulation^9,30^, MHC II is subject to regulation of intracellular protein trafficking, including translocation to the cell surface, endocytosis, and degradation^10^. Therefore, we determined whether NRF2 regulation affects total intracellular MHC II protein expression or only surface expression at baseline or after stimulation. To do so, we took advantage of dual-MHC II antibody labeling before and after permeabilization that allowed us to detect both surface and internal protein expression by standard and imaging flow cytometry (**Fig 3C-E**). We found that internal expression of MHC II in NRF2^-/-^ mexAMs was significantly increased 1.686-fold over WT controls at baseline (**Fig 3C**), a difference retained following LPS stimulation, but not IFNɣ stimulation (**Fig 3D, E**). Given that IFNɣ is such a strong promoter of *Ciita* and MHC II expression^9,10,30^, this data suggests that although NRF2 inhibits internal protein expression at baseline and after a weak stimulus, such as LPS, regulation of internal MHC II levels can be superseded by a strong stimulus, such as IFNɣ. Overall, our results demonstrate that NRF2 regulates MHC II surface expression and also its intracellular levels in AMs and mexAMs. Although these results do not rule out an effect of NRF2 on MHC II intracellular trafficking, per se, they suggest that the primary regulation by NRF2 is at the level of MHC II gene expression and total protein levels.

**Figure 3:**
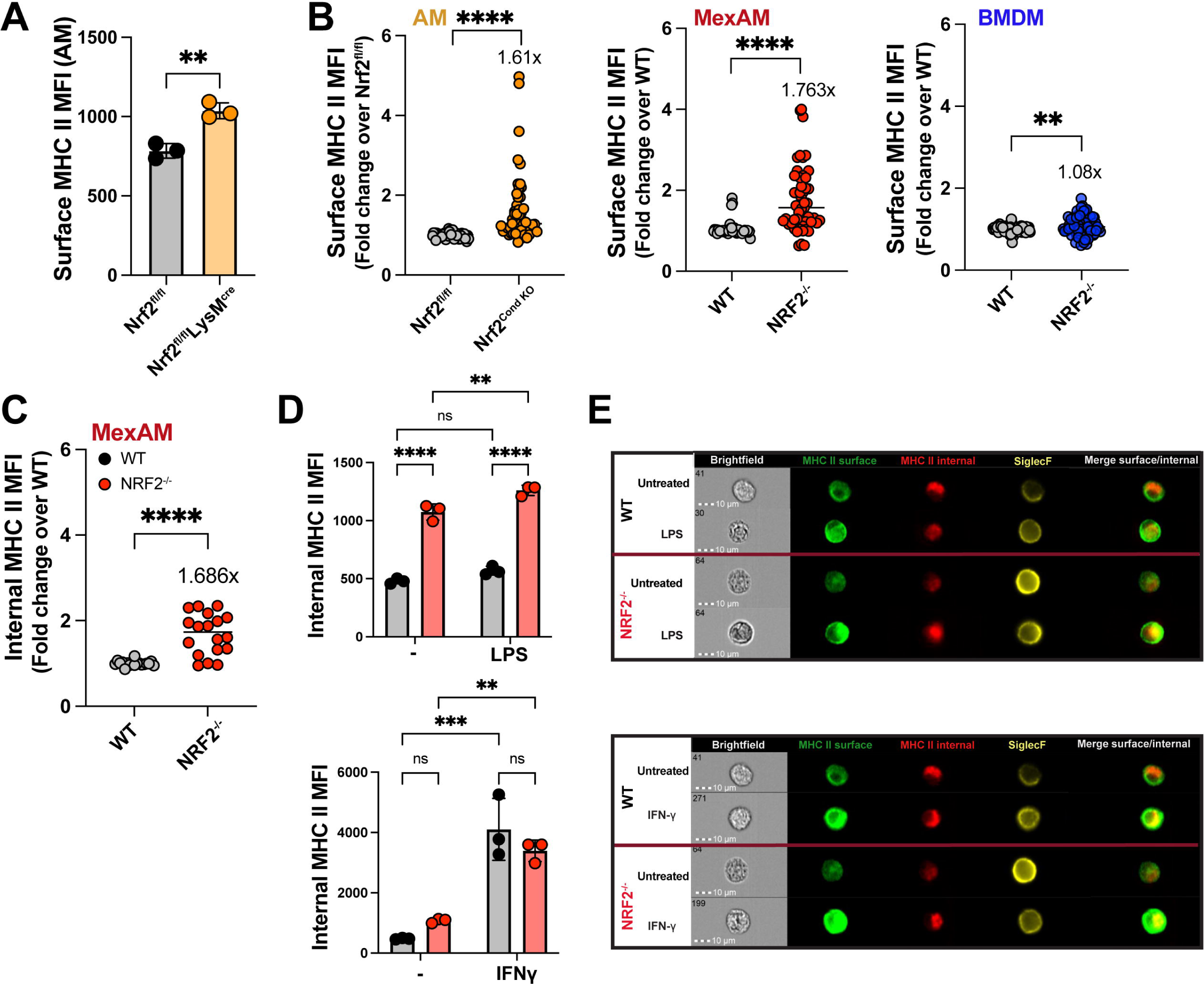
NRF2 inhibits surface and internal MHC II protein levels in AMs, but not BMDMs. **A)** Surface MHC II MFI in Nrf2^fl/fl^ and Nrf2^fl/fl^LysM^cre^ AMs. **B)** Surface MHC II MFI in Nrf2^condKO^ AMs, NRF2^-/-^ mexAMs, and NRF2^-/-^ BMDMs, normalized to WT. **C)** Internal MHC II MFI in WT and NRF2^-/-^ mexAMs, normalized to WT. **D, E)** WT and NRF2^-/-^ mexAMs treated for 24 hours with either LPS (100 ng/mL) (top) or IFNγ (20 ng/mL) (bottom). Internal MHC II MFI **(D),** surface and internal MHC II staining **(E)**. **A)** Representative of 19 independent experiments **B)** Compiled from more than 19 independent experiments. Each dot represents a technical replicate. **C-D)** Representative of 3 or more independent experiments. **A,B)** Unpaired Student’s t-test. **C-D)** Two-way ANOVA with Tukey post-test.*p <0.05, **p< 0.01, ***p< 0.001, ****p< 0.0001.

### NRF2 repression of AM MHC II is retained after PAMP and cytokine stimulation

Given the many signals known to up-regulate surface expression of MHC II on innate cells^10^, we next investigated the influence of NRF2 on MHC II expression following exposure to various PAMPs and Type I and II IFNs. We exposed WT and NRF2^-/-^ mexAMs to ligands of different Toll-like receptors, including LPS (TLR4 ligand), Pam3Cys (TLR1/2 ligand), CpG (TLR9 ligand), and R848 (TLR7/8) in addition to 2’3’-cGAMP, a ligand of STING. These PAMPs were chosen to represent innate signals from both bacterial and viral infections, including many associated with Mtb infection^31^ **(Fig 4A)**. We found that NRF2^-/-^ mexAMs consistently exhibited higher MHC II expression compared to WT mexAMs, across these varied stimuli. We next evaluated a role for NRF2 regulation of MHC II following IFNɣ and IFNβ stimulation **(Fig 4B**). IFNβ can promote MHC II expression^32^ but is typically associated with TB disease progression rather than protection^33–36^. IFNɣ is a robust inducer of MHC II and activates macrophages that leads to restriction of Mtb growth and resistance to Mtb^37–39^. Despite no difference in internal MHC II expression between WT and NRF2^-/-^ mexAMs after IFNɣ stimulation (**Fig 3D**), NRF2^-/-^ mexAMs had higher surface MHC II expression than WT mexAMs after IFNɣ stimulation **(Fig 4B**). These results indicate that NRF2 still regulates surface MHC II expression after IFNɣ stimulation, despite well-documented IFNɣ regulation of the CIITA pathway^40^. In contrast, WT and NRF2^-/-^ mexAMs expressed equivalent levels of surface MHC II after IFNβ stimulation **(Fig 4B**). Primary AMs lacking NRF2 also retained higher MHC II levels than WT controls after both LPS and IFNɣ stimulation **(Fig 4C)**.

**Figure 4:**
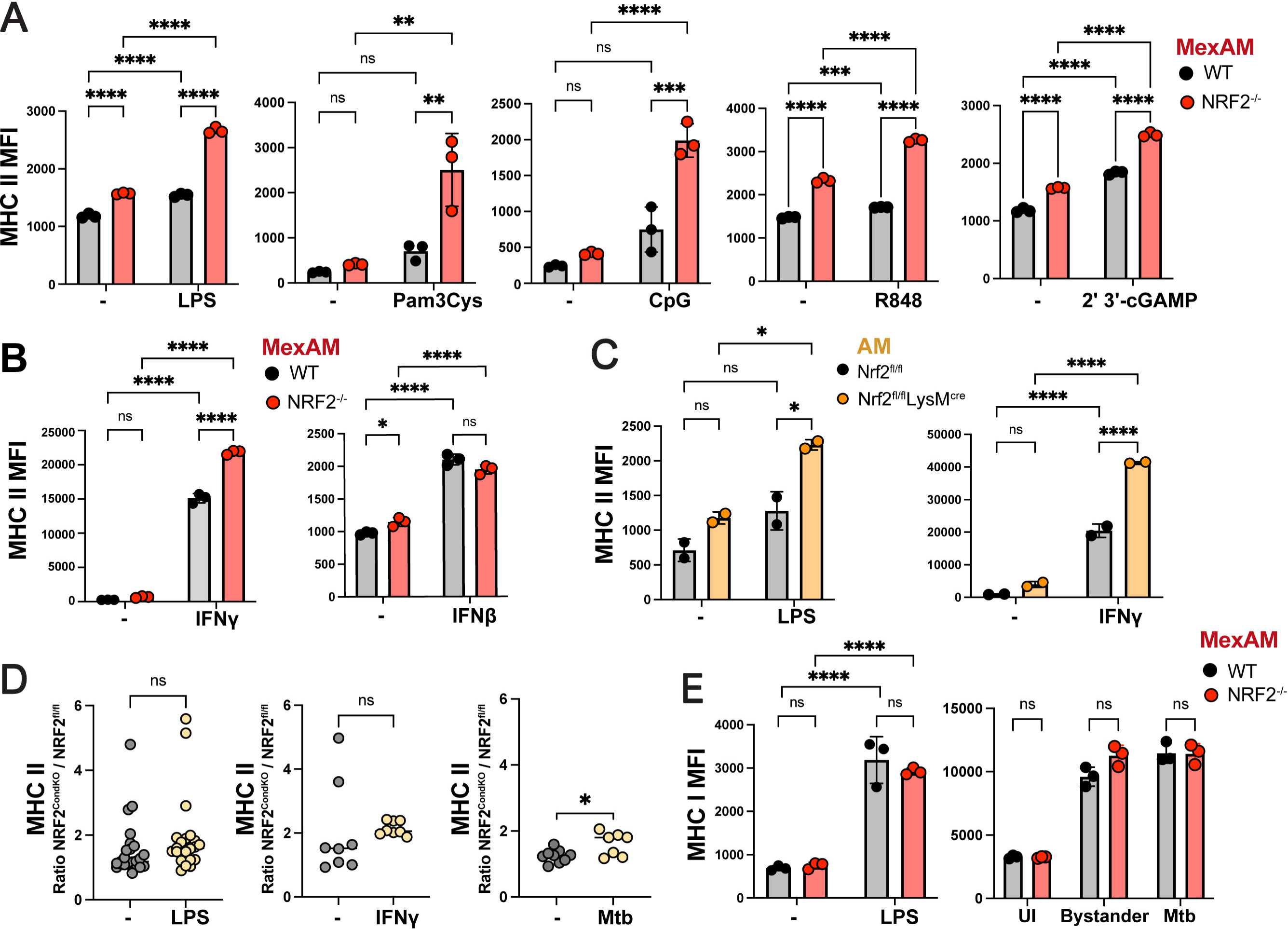
NRF2 inhibition of AM MHC II is retained after PAMP and IFNγ stimulation. **A)** MHC II MFI for WT and NRF2^-/-^ mexAMs stimulated with LPS, Pam3Cys, CpG, R848, or 2’3’-cGAMP for 24 hours. **B)** MHC II MFI for WT and NRF2^-/-^ mexAMs stimulated with IFNγ or IFNβ for 24 hours. **C)** MHC II MFI for Nrf2^fl/fl^ and Nrf2^fl/fl^ LysM^cre^ AMs, LPS or IFNγ stimulation. **D)** MHC II MFI ratio for Nrf2^condKO^ to Nrf2^fl/fl^ AMs, LPS, IFNγ, or Mtb stimulation. **E)** MHC I MFI for WT and NRF2^-/-^ mexAMs after LPS stimulation or Mtb infection. **A-C, E)** Representative of 3 or more independent experiments. **D)** Compiled from 3+ independent experiments. **A-C, E)** Two-way ANOVA with Tukey post-test. **D)** Unpaired Student’s t-test. *p<0.05, **p< 0.01, ***p< 0.001, ****p< 0.0001.

Considering that MHC II expression was higher in NRF2^-/-^ AMs at baseline and after stimulation, we next determined whether the difference in MHC II levels between WT and NRF2^-/-^ AMs was further enhanced by stimulation, retained at baseline levels, or diminished. We compared the ratio of MHC II expression in Nrf2^CondKO^ to Nrf2^fl/fl^ AMs in untreated versus LPS, IFNɣ, and Mtb-infection conditions (**Fig 4D**). The expression ratios for LPS or IFNɣ conditions were equivalent to the ratio of untreated AMs, and the Mtb-infection condition was slightly higher. These results indicate that not only are the baseline levels of MHC II expression in AMs regulated by NRF2, but that NRF2 maintains a similar impact on MHC II expression after stimulation. Overall, we find that AMs lacking NRF2 retain a higher expression of MHC II than WT AMs across diverse PAMP and IFNɣ stimulations. Our data strongly indicate that NRF2 not only regulates the baseline levels of MHC II expression in AMs and mexAMs, but that this effect also impedes MHC II expression upon microbial stimulation.

To determine the effect of NRF2 on MHC II in inflammatory macrophages, we repeated these PAMP stimulations in WT and NRF2^-/-^ BMDMs in the presence or absence of the Nrf2 agonist, L-sulf. For LPS stimulation, we observed a significant decrease in MHC II expression following L-sulf treatment in WT, but not NRF2^-/-^, BMDMs, suggesting that exogenous NRF2 activation can impede LPS-induced MHC II expression in BMDMs (**Fig S4A**). In contrast, MHC II induction by 2’3’-cGAMP was not inhibited by L-sulf treatment (**Fig S4A**). To compare the effect of NRF2 activation across stimulation conditions, we calculated the percent inhibition of MHC II by L-sulf in BMDMs across 5 different PAMPs. We found that MHC II up-regulation by LPS, Pam3Cys, CpG and R848, but not 2’3’-cGAMP, was significantly inhibited by L-sulf in a NRF2-dependent manner (**Fig S4B**). Overall, these results show that NRF2 modulates MHC II expression in inflammatory macrophages under most stimulation conditions, when NRF2 is activated.

We also measured MHC II expression in WT and NRF2^-/-^ bone marrow-derived dendritic cells (BMDCs) after LPS stimulation in the presence of absence of L-sulf. As expected, MHC II expression was much higher in BMDCs than in macrophages. However, we observed a similar increase of MHC II in WT and NRF2^-/-^ cells upon LPS stimulation, with very minimal effect of NRF2 activation by L-sulf treatment (**Fig S4C**). To determine whether NRF2 impacts all antigen presentation pathways or only MHC II, we measured MHC I in WT and NRF2^-/-^ mexAMs following LPS stimulation and Mtb infection. As expected, both LPS and Mtb led to an increase in MHC I expression (**Fig 4E**). However, there was no differences in MHC I expression between WT and NRF2^-/-^ cells, indicating that MHC I expression is not regulated by NRF2. Overall, NRF2 inhibition of antigen presentation affects MHC II but not MHC I, and is specific to macrophages, and not DCs.

### NRF2 inhibition of MHC II in AMs inhibits antigen-specific CD4^+^ T cell activation

Previous studies have found that human AMs are not effective at presenting antigens to T cells^41–44^, and in some cases can induce T cell unresponsiveness^45^. To determine whether NRF2 inhibition of AM MHC II impedes their ability to activate CD4^+^ T cells, we established an *in vitro* co-culture system based on previous studies^46^. MexAMs or primary AMs were loaded with different concentrations of ESAT6_3-17_ peptide (0.1-10µM) and co-cultured with C7 TCR transgenic CD4^+^ Th1 cells (hereafter, C7 T cells), which express a TCR specific for the ESAT6_3-17_ Mtb peptide, at a ratio of 1:1 **(Fig 5A, S5A)**. When cultured with WT macrophages loaded with increasing concentration of peptide, C7 T cells produced increasing amounts of IFNɣ. C7 T cells produced significantly more IFNɣ when cultured with peptide-pulsed NRF2^-/-^ mexAMs or AMs than WT cells. A similar trend was observed in the BMDM C7 T cells co-culture system under exogenous NRF2 activation, where co-culture with NRF2^-/-^ BMDMs led to significantly higher IFNɣ production by C7 T cells than WT BMDMs in the presence of L-sulf **(Fig S5B)**. Next, we cultured C7 T cells with H37Rv-infected WT or NRF2^-/-^ mexAMs. Again, C7 T cells produced more IFNɣ when cultured with Mtb-infected NRF2^-/-^ mexAMs compared to infected WT mexAMs **(Fig 5B)**. The expression of MHC II and CD86, a T cell co-stimulatory molecule and a marker of macrophage activation, was measured on WT and NRF2^-/-^ Mtb-infected macrophages 24 hours after culture with C7 T cells. We observed significantly higher levels of MHC II and CD86 on NRF2^-/-^ mexAMs compared to WT mexAMs (**Fig 5C**). In parallel, we examined C7 T cell expression of CD69, a marker of T cell activation after culture with mexAMs. CD69 expression was higher on C7 T cells cultured with Mtb-infected NRF2^-/-^ mexAMs compared to WT mexAMs (**Fig 5D**). We also confirmed that the up-regulation of CD86 on Mtb-infected WT and NRF2^-/-^ mexAMs was dependent on the presence of T cells **(Fig 5E)**. To confirm that differences in IFNɣ production or activation markers were not due to any difference in initial Mtb uptake between WT and NRF2^-/-^ mexAMs, we measured Colony Forming Units (CFU) as a measurement of bacterial burden on day 1 of infection, a timepoint before C7 T cells would provide any protective effects against bacterial growth. There was no difference in the bacterial burden of WT and NRF2^-/-^ mexAMs **(Fig 5F)**. These results demonstrate a critical role for NRF2-mediated regulation of macrophage MHC II in T cell activation during Mtb infection and suggest that NRF2 can limit the ability of infected macrophages to promote effective adaptive immune responses.

**Figure 5:**
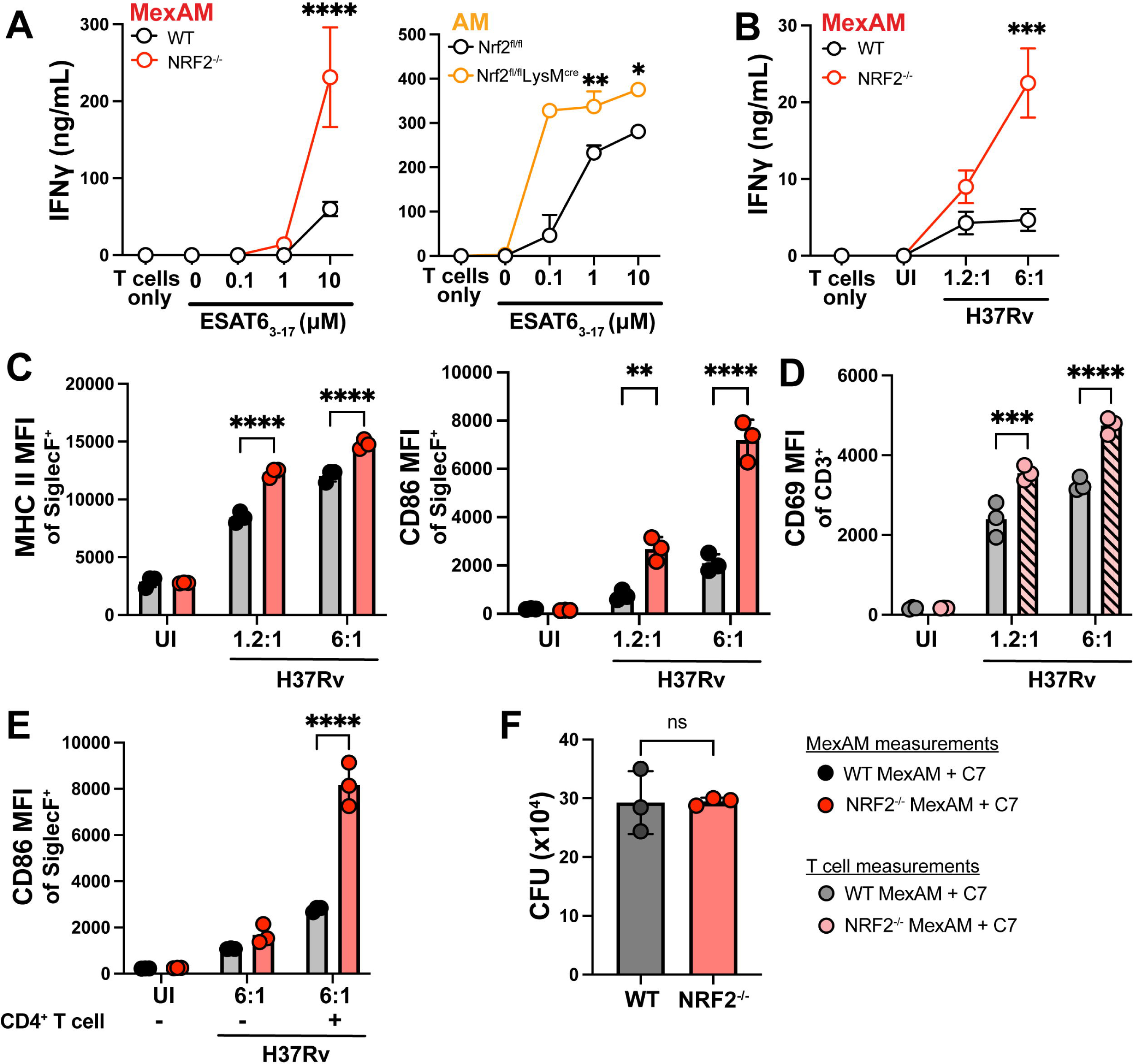
Absence of NRF2 in AMs and mexAMs enhances antigen-specific CD4^+^ T cell activation. **A)** IFNγ production following 72 hour co-culture of C7 T cells and WT or NRF2^-/-^ mexAMs and WT or Nrf2^cond^ ^KO^ primary AMs loaded with 0.1-10μM ESAT6_3-17_ peptide. **B)** IFNγ production following 72 hour co-culture of C7 T cells and WT or NRF2^-/-^ mexAMs infected with mCherry-H37Rv (MOI 1.2:1 and MOI 6:1). **C)** MHC II and CD86 MFI of Siglec F^+^ cells, 24 hours following co-culture. **D)** CD69 MFI of CD3^+^ cells 24 hours following co-culture. **E)** CD86 MFI of Siglec F^+^ cells, 24 hours following co-culture, with or without CD4^+^ T cells. **F)** CFU of Mtb-infected WT or NRF2^-/-^ mexAMs co-cultured with CD4^+^ T cells 1 dpi. **A)** Representative of 3 independent experiments (mexAMs) or 1 independent experiment (primary AMs). **B-F)** Representative of 3 independent experiments. **A-E)** Two-way ANOVA with Tukey post-test. **F)** Unpaired Student’s t-test. *p <0.05, **p< 0.01, ***p< 0.001, ****p< 0.0001.

## DISCUSSION

AMs occupy a unique niche within the lung alveolar spaces and are critical in the early response to Mtb infection^2,3,47^. Despite their early sentinel role, AMs are hyporesponsive during the initial phases of Mtb infection^3,8,48–50^. In this context, NRF2, a central regulator of cellular oxidative stress responses, serves as a critical modulator of AMs during Mtb infection. We previously showed that as early as 12 hours after infection AMs mount a cell-protective response, rather than proinflammatory response, which is driven by NRF2^3^. NRF2 is known to regulate many processes including antioxidant production, cell death, and iron sequestration, but not all of its roles are yet appreciated^16–18,20,51,52^

By specifically examining AMs *in vitro* and *in vivo*, our study reveals that NRF2 inhibits MHC II expression on Mtb-infected AMs and on AMs under various other stimulation conditions. Through examination of gene expression and protein localization, we determined how NRF2 regulates MHC II in AMs. We observed that NRF2 limits expression of *Ciita*, a critical master regulator of the MHC II locus, as well as inhibition of MHC II genes (i.e. *H2-Ab1)* and total MHC II protein expression. In contrast, the expression of MHC I is not affected by NRF2. NRF2-mediated inhibition of MHC II expression on AMs is evident at baseline and is retained after stimulation with innate immune activators and with IFNɣ. NRF2 regulation of MHC II at the level of *Ciita* transcription was unexpected. Our previous study detected 583 genes with NRF2 binding motifs by HOMER and 3,975 NRF2 ChIP-seq binding peaks in macrophages (GSE75175) but found no evidence for NRF2 binding at the *Ciita* promoter region (defined as 2,000 nucleotides upstream to 1,000 nucleotides downstream from the gene start site)^3^. Therefore, we hypothesize that regulation of *Ciita* by NRF2 may be indirect. Interestingly, a recent study demonstrated that KEAP1, an inhibitor of NRF2, promotes IFNɣ-induced *Ciita* and MHC II expression in HeLa cells through inhibition of HDAC activity^53^. However, through experiments that co-depleted KEAP1 and NRF2, the authors concluded that the KEAP1 effect was independent of NRF2, a distinction from our results that could be due to differences in cell type.

These effects are specific to AMs; NRF2^-/-^ AMs at baseline express higher levels of *Ciita* and *H2-Ab1* than WT AMs, while NRF2^-/-^ BMDMs do not express higher levels than WT BMDMs. This cell-specific effect is likely a result of the difference in baseline NRF2 activity between AM and BMDMs. While Mtb-infected AMs significantly up-regulate the NRF2 transcriptional program over levels of naïve AMs *in vivo*, our results indicate that AMs already have relatively high NRF2 activity at baseline. We detected NRF2 accumulation by microscopy in AMs, but not BMDMs, after LPS stimulation (**Fig S2A, B**), as well as higher expression levels of NRF2 target gene, *Nqo1* in AMs (**Fig 2A**). Additionally, we found that when BMDMs were treated with a NRF2 agonist, a difference in MHC II expression between WT and NRF2^-/-^ cells was revealed. Interestingly, WT and NRF2^-/-^ BMDCs also expressed equivalent levels of MHC II at baseline, yet L-sulf treatment did not yield differences in MHC II expression. The fact that exogenous NRF2 activation affects MHC II expression in macrophages but not DCs highlights the fundamental differences in MHC II regulation across cell types. In further support of an AM-specific effect, our data demonstrate that, in Mtb-infected lungs, NRF2 inhibits MHC II expression on AMs, but not on pulmonary MDMs, DCs or PMNs.

Overall, our results suggest that macrophages with high NRF2 activity will have impeded MHC II presentation. We surmise that because AMs have high NRF2 activity at baseline, NRF2 inhibits MHC II under many conditions. However, NRF2 regulation of MHC II may be restricted to specific conditions for other macrophage subsets. For example, recent single-cell RNA-sequencing studies have identified CD38^-^, Nos2^-^ and iron-associated subsets of interstitial macrophages (IMs) with high expression of NRF2 and NRF2-associated oxidative stress response genes between 2-4 weeks following Mtb infection^48,49,54^. Based on our results in AMs, we would predict these IM populations with high NRF2 expression may also have impaired antigen presentation.

Activation of CD4^+^ T cells is a critical step for an effective host response to Mtb^55–59^. AMs serve as primary targets for Mtb during the earliest stages of infection, but do not migrate out of the lung. Instead, Mtb-infected monocytes or DCs eventually migrate to the lung-draining lymph node and transfer bacterial antigens to uninfected DCs in the lymph node for initial priming of CD4+ and CD8^+^ T cells^60–62,63^. Primed CD4^+^ T cells are recruited to the pulmonary infection site, where their production of IFNɣ and other effector functions are crucial for Mtb control^64–68^. CD4^+^ T cell help also prevents CD8^+^ T cell exhaustion and preserves their antibacterial function, which promotes host control of Mtb infection^56^. While it is well-appreciated that direct contact with infected cells is required for the cytotoxic function of CD8^+^ T cells, surprisingly effector CD4^+^ T cells must also directly recognize MHC II-expressing infected cells for effective Mtb control^56^. This requirement was demonstrated by Srivastava et al, who used mixed bone marrow chimeric mice to show that MHC II^-/-^ DCs and recruited macrophages contained greater numbers of bacteria than MHC II^+/+^ cells, and this difference was lost after CD4^+^ T cells depletion^59^. These results highlight how increased NRF2 expression could impede CD4+ T cell responses in the lung by constraining MHC II expression by infected macrophages.

NRF2 expression is only one of several mechanisms that can limit pulmonary T cell responses during Mtb infection. A mismatch between the antigens presented by uninfected DCs during T cell priming in the draining lymph node and the antigens presented by infected macrophages in the lung can also lead to poor T cell recognition of Mtb-infected macrophages^69^. Furthermore, virulent Mtb can inhibit CIITA and MHC II presentation^64^.

The consequences of NRF2-mediated inhibition of MHC II are critical to CD4^+^ T cell responses and macrophage activation through crosstalk. In our co-culture system, the absence of NRF2 in AMs significantly increased IFNɣ production by CD4^+^ T cells and enhanced activation of both cell types, based on increased expression of CD69 on T cells and CD86 on AMs. IFNɣ activates macrophages, which enhances phagocytosis, induces nitric oxide, and promotes lysosomal fusion and degradation of phagosomal contents, antimicrobial activities that restrict Mtb growth and promote bacterial clearance^70–74^. Effective CD4^+^ T cell responses are crucial to coordinate granuloma formation, which are structured aggregates that encapsulate infected macrophages and limit bacterial dispersal within the lung^75,76^. Several recent studies find heterogeneity in the permissiveness of different myeloid cells in the lung after the initiation of the adaptive response^48,50,75^. Whether NRF2 inhibition of MHC II contributes to these dynamics is an important question for further consideration.

Lastly, the impact of NRF2 on macrophage antigen presentation may extend beyond effector CD4^+^ T cells. Memory CD4^+^ T cells also require antigen presentation by MHC II for homeostatic maintenance^77,78^. Therefore, both recognition of infected macrophages by effector CD4^+^ T cells in the lung during active infection, and the persistence of long-lived memory CD4^+^ T cells following vaccination or infection could be impeded when T cells interact with AMs or IMs with high NRF2 activity.

In conclusion, understanding the impact of NRF2 on AM immune responses and antigen presentation provides insight into the complex interplay between innate and adaptive immunity during Mtb infection. Targeting NRF2 signaling pathways in AMs represents a promising avenue for therapeutic interventions aimed at enhancing host immune defenses into limit bacterial growth^20,79^. Blockade of NRF2 could promote CD4^+^ T cell production of IFNɣ, a central focus of current TB vaccine candidates^69^. However, separating the inhibition of antigen presentation from other functions of NRF2 that are critical to dampen tissue inflammation and limit lung pathology will be a challenge^80,81^. Therefore, future research efforts should focus on elucidating the precise molecular mechanisms underlying NRF2-mediated immune modulation, with implications for developing novel immunotherapeutic strategies against Mtb and other infectious diseases.

### Limitations of the study

We demonstrate the importance of NRF2 in regulating antigen presentation by AMs. While we show that NRF2 regulates *Ciita* and *H2-Ab1* gene expression and MHC II protein expression in AMs, without affecting MHC I, the molecular mechanism that links NRF2 and MHC II gene expression should be the subject of future studies, especially given that the *Ciita* promoter does not contain a known NRF2-binding site. Our use of ESAT6_3-17_-specific C7 transgenic CD4^+^ T cells show how NRF2 modulation of the MHC II pathway affects T cell activation. Examining a broader repertoire of T cell responses, including the Ag85b_240-254_-specific P25 transgenic CD4^+^ T cells^46^, as well as *in vivo* primary T cell responses, would generalize how NRF2 affects T cell recognition. Due to the limitations of the NRF2 knockout and conditional knockout mouse strains with potential effects on non-AM cells^82,83^ and the feasibility of isolating sufficient numbers of cells, primary AMs weren’t used in all experiments. Instead, we chose to use mexAMs at low passage numbers, as an alternative tool. Another question that remains unanswered is when during infection NRF2-regulated MHC II expression and reduced T cell activation impacts the endogenous host response and bacterial clearance. Further studies are needed to investigate the downstream impact of NRF2 on MHC II in macrophages and its role in both effector and memory CD4^+^ T cell responses. Finally, future studies should investigate the role of NRF2-regulated MHC II expression on AMs following vaccination, especially considering the crucial role of MHC II expression in maintaining memory CD4^+^ T cells^77,78^, the importance of airway memory CD4^+^ T cells in protection from Mtb^69^, and AM remodeling in the airway after exposure to *M. bovis* BCG vaccination^84^. Addressing these key factors would provide valuable insights for therapeutic and vaccine development.

## MATERIALS AND METHODS

### Mice

WT C57BL/6J and Nrf2^-/-^ (B6.129X1-Nfe2l2^tm1Ywk^/J) mice were obtained from the Jackson Laboratory (Bar Harbor, ME). NRF2 conditional KO mice, Nrf2^floxed^ (C57BL/6-*Nfe2l2t^m1.1Sred^*/SbisJ), CD11c^cre^ [B6.Cg-Tg(Itgax-cre)1-1Reiz/J], and LysM^cre^ [B6.129P2-*Lyz2^tm1(cre)Ifo^*/J] mice were also purchased from the Jackson Laboratory and crossed to generate Nrf2^CD11c^ and Nrf2^LysM^ transgenic lines. The animals were housed under specific pathogen-free conditions at the University of Massachusetts Amherst vivarium, and all experiments were conducted in compliance with the Institutional Animal Care and Use Committee guidelines. Male and female mice aged 6 to 12 weeks were utilized for the experiments.

### *Mycobacterium tuberculosis* aerosol infections and lung cell isolation

Aerosol infections were performed with H37Rv-mCherry at standard dose (∼100 CFU) infection. Mice were enclosed in an aerosol infection chamber (Glas-Col) and frozen stocks of bacteria were thawed and placed inside the associated nebulizer. To determine the infectious dose, three mice in each infection were sacrificed one day later and lung homogenates were plated onto 7H10 plates for CFU enumeration. Analysis of experimental samples was performed on either 12 or 14 days post-infection. Lungs were removed and single-cell suspensions were prepared by Liberase (Roche) digestion containing DNase I (Sigma-Aldrich) for 30 minutes at 37°C. Mechanical disruption was performed with the Gentle-MACS dissociator (Miltenyi Biotec), followed by double filtering through a 70 μM cell strainer and removal of red blood cells using ACK lysis buffer (Gibco). Cells were resuspended in FACS buffer (PBS, 1% FBS, 0.1% NaN_3_) prior to staining for flow cytometry.

### Alveolar macrophage isolation

Bronchoalveolar lavage (BAL) was carried out on mice from WT and Nrf2^LysM^ or Nrf2^CD11c^ after euthanasia, as previously described^85^. Briefly, the trachea was exposed, and an incision was made using Vannas Micro Scissors (VWR). A 20-gauge, 1-inch intravenous catheter (McKesson) connected to a 1 mL syringe was used to instill 1 mL of phosphate-buffered saline (PBS) into the lungs. The PBS was gently flushed and aspirated three to four times, and the recovered fluid was immediately placed on ice. This lavage process was repeated three to four additional times to collect the maximum number of cells. The collected samples were filtered through a 70µm strainer, and the cells were pelleted by centrifugation. For antibody staining experiments, the cell pellet was resuspended in FACS buffer solution. Alternatively, for cell culture assays, the cells were seeded at a density of 1 × 10^5^ cells per well in a 96-well plate using complete RPMI medium, with RPMI (Millipore Sigma, 11875-119), 10% FBS (Biowest), 1% L-glutamine (2 μM, Gibco) and 1% penicillin streptomycin (100 U/mL, Gibco)

### Bone marrow-derived macrophage isolation

Bone marrow-derived macrophages (BMDMs) were obtained by harvesting femurs from either WT or NRF2^-/-^ mice strains. The cells collected from the bone marrow were cultured in complete RPMI medium supplemented with 50 ng/mL recombinant human macrophage colony-stimulating factor (M-CSF, Peprotech) for a period of 6 days to promote differentiation into macrophages, BMDM media was changed on day 4 of culture

### Murine ex vivo alveolar macrophage generation

Primary AMs were isolated by BAL from WT and NRF2^-/-^ mice as described above. AMs were cultured in complete media containing RPMI media, 10% fetal bovine serum (VWR), 2 mM L-glutamine (Invitrogen), 100 U/mL penicillin-streptomycin (Invitrogen), and 30 ng/mL GM-CSF (PeproTech), 10ng/ml TGF-β (PeproTech), and 1uM rosiglitazone (Sigma-Aldrich).

### Macrophage stimulation

Macrophages from BMDMs, mexAMs, or AMs were stimulated with PAMPs for a range from 4 hours to 24 hours depending on the analysis, including flow cytometry and ELISA (24 hours), or RT-qPCR (4 hours). PAMPs include Lipopolysaccharide (LPS) R595 (List labs), Pam3Cys (Pam3CSK4) (Invivogen), 2’3’-cGAMP (Invivogen), Resiquimod (R848) (Sigma-Aldrich), CpG (Invivogen), Recombinant Mouse IFN-β1 (BioLegend), Recombinant Mouse IFNƔ (Peprotech)

### Flow cytometry staining

Cells were resuspended in FACS buffer (PBS with 1% FBS and 0.01% NaN_3_) and Fc receptors were blocked with anti-CD16/32 (Biolegend). Cell viability was assessed with Zombie Violet dye (BioLegend). Surface staining included antibodies specific for murine, including Siglec F (clone S17007L), CD11c (clone N148), F4/80 (clone BM8). The following antibodies were used for lung digestion biomarkers I-A/I-E, CD86, CD69, CD11b, CD64, CD45, CD3, CD19, CD11c, Ly6G, and Ly6C (reagents from BioLegend unless otherwise noted). For lung digestion, cell numbers were enumerated using Polybead Polystyrene 15 μm Microspheres (Polysciences). Cells were then fixed in 2% PFA (Electron Microscopy Sciences) at room temperature. Samples were then washed with FACS buffer and resuspended in FACS buffer for immediate use or stored overnight at 4°C. Samples were analyzed on a 3- or 5-laser LSRFortessa flow cytometer (BD Bioscience) and FlowJo software (Tree Star, Inc.).

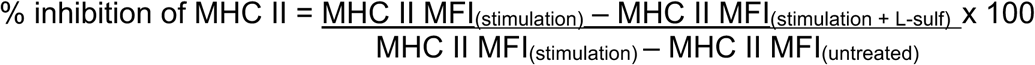

### Internal and surface MHC II staining

Similar to standard staining for flow cytometry, as described above, cells were collected into FACS tubes and subsequently stained with Fc block (Biolegend), Live/Dead stain (Zombie Violet Fixable Viability Kit, Biolegend) and characteristic cell surface markers (AMs or mexAMs, Siglec-F (Biolegend),; BMDMs F4/80 (clone BM8, Biolegend), and I-A/I-E antibody (Biolegend) for surface MHC II marker. Cells were resuspended in 200 μL of Cyto-Fast™ Fix/Perm Buffer (Biolegend) and incubated for 20 minutes at room temperature and washed with Cyto-Fast™ Perm Wash (Biolegend). Another I-A/I-E antibody in a different channel was diluted 1:100 in Cyto-Fast™ Perm Wash and added to the cells for 20 minutes. Cells were then washed and fixed with 2% PFA at room temperature. Samples were then washed with FACS buffer and resuspended in FACS buffer for immediate use or stored overnight at 4°C. Samples were analyzed on a 3- or 5-laser LSRFortessa flow cytometer (BD Bioscience) and FlowJo software (Tree Star, Inc.).

### Enzyme-linked immunosorbent assay (ELISA)

Cells were plated at 0.5 - 1×10^5^ density in either 96 or 24 well plates, respectively, followed by different stimulation. Supernatants were collected 24 hours after macrophage stimulation, or 72 hours after macrophage - C7 T cells co-culture. Murine IFNɣ DuoSet ELISA were performed following manufacturer’s instructions (R&D Systems)

### RNA isolation and quantitative reverse transcription PCR (RT-qPCR)

Primary cells or *ex vivo* cultured cells were plated at a 0.5 - 1×10^5^ density in either 96 or 24 well plates, respectively. RNA isolation was performed using TRIzol (Invitrogen), followed by two sequential chloroform extractions, addition of GlycoBlue (Invitrogen), isopropanol precipitation, and 2 washes of 70% ethanol, with final resuspension in RNase free distilled water. RNA was quantified using the Biodrop duo (Biochrom). RNA amounts of 1ug per sample were converted to complementary DNA (cDNA) then amplified using RNA to cDNA EcoDry Premix (TaKaRa). TaqMan primer probes (IDT) were used for genes of interest: *Nqo1*, *Ciita* (ThermoFisher), *H2-Ab1* (IDT). TaqMan Fast Universal PCR Master Mix (Applied Biosystems) was used for the RT-qPCR reaction, performed by BioRad CFX Opus 96 RT-qPCR machine. All the generated data were then normalized relative to *Gapdh* or *Ef1a* housekeeping genes in each sample.

### Expansion of transgenic C7 CD4^+^ T cell line

C7 CD4^+^ T cells were a gift from Dr. Samuel Behar from UMass Chan, Worcester, MA. C7 TCR transgenic mice were originally generated by Dr. Eric Palmer^86^. C7 CD4^+^ T cells were stimulated *in vitro* with irradiated splenocytes pulsed with the ESAT-6_3−17_ peptide, in complete RPMI media containing 10% fetal bovine serum (VWR), 2 mM L-glutamine (Invitrogen), 100 U/mL penicillin-streptomycin (Invitrogen), 2-Mercaptoethanol (Gibco) and 20 U/mL recombinant murine IL-2 (Peprotech). After the initial stimulation, T cells were cultured in complete media containing recombinant murine IL-2 along with 0.01 μg/mL recombinant murine IL-7 (Peprotech). Cells were split every two days for 3–4 divisions and then rested for two to three weeks before use.

### T cell-macrophage co-culture

The following synthetic peptide epitope was used as an antigen for the T cell-macrophage co-culture : ESAT-6_3−17_ (EQQWNFAGIEAAASA). The peptides were synthesized by Genscript. The macrophages were pulsed with 0.1-10 μM of the peptide for 1 hour. After incubation, the macrophages were washed 3 to 5 times and then resuspended in complete RPMI 1640 media prior to addition of the CD4^+^ T cells. For *in vitro* infections, macrophages were infected with mCherry-H37Rv at 1.2:1 or 6:1 MOIs. Two hours after the infection, infected macrophages were washed 3 times with RPMI and C7 T cells were added with a 1:1 ratio of macrophages:T cells

### Mtb *in vitro* infection

mCherry-H37Rv was cultured from frozen stock in 7H9 media for 2 days to an OD_600_ of 0.1-0.3. Bacteria was added for two hours and then macrophage cultures were washed three times to remove extracellular bacteria. The final concentration was calculated based on strain titer to produce an effective MOI of 1.2:1, 2.4:1, and 6:1 after two hours of incubation and washing.

### Immunofluorescence microscopy

BMDMs and mexAMs from WT and NRF2^-/-^ strains were seeded at 100,000 cells per well in an 8-well chamber slides (Thermo Fisher Lab-Tek^Ⓡ^II Chamber Slide^TM^). Cells were incubated in a 5% CO_2_ incubator at 37 °C. To identify NRF2 activation, cells were treated with L-sulforaphane overnight. Cells were washed with PBS followed by fixation with 4% paraformaldehyde for 15 minutes at room temperature. Cells were incubated in blocking buffer (containing 1X PBS, 5% normal Goat serum, 0.3% Triton X-100) for 1 hour with anti-Nrf2 (D1Z9) XP Rabbit Antibody (Cell signaling) or isotype control antibody Rabbit (DA1E) Antibody IgG (Cell signaling) and incubated at 4°C overnight. The next day, after washing cells three times with PBS, secondary antibody Anti-rabbit IgG Fab2 AF488 (Cell Signaling) was added, followed by 3 washes in PBS and 2 washes in 3% BSA. The nuclei were stained with Hoechst (DAPI 20mM, Thermo Fisher) for 15 minutes, followed by 2 washes in 3% BSA and two washes in PBS. Slides were then mounted and allowed to air-dry at room temperature, avoiding direct light exposure. The next day, slides were imaged using the Nikon Ti2 stand with A1HD(1024) Resonant Scanning Confocal.

### Statistical analysis

Statistical analysis was performed using Graphpad Prism 9. P-values were calculated using unpaired t-test (2 conditions), one-way ANOVA or two-way ANOVA with Tukey post-test (3+ conditions), or as indicated in the figure legends. P-values were indicated as follow: \**P* < 0.05, \*\**P* < 0.01, \*\*\**P* < 0.001, \*\*\*\**P* < 0.0001.

## Acknowledgments

We thank the Animal Care Staff at the University of Massachusetts Amherst. We thank Amy Burnside at the Flow Cytometry Core, and James Chamber at the Light Microscopy at the University of Massachusetts Amherst. We thank members of the Rothchild and Behar labs for helpful discussions.

## Author contributions

Conceptualization: LKP, ACR

Methodology: LKP, TS, ACR

Investigation: LKP, MMC, PNL, DD, AT, TS, ACR

Visualization: LKP, ACR

Data curation: LKP, ACR

Formal analysis: LKP, ACR

Project administration: ACR

Funding acquisition: SMB, ACR

Supervision: SMB, ACR

Writing – original draft: LKP, ACR

Writing – review & editing: LKP, MMC, PNL, DD, TS, SMB, ACR

## SUPPLEMENTARY FIGURE LEGENDS

**Figure S1: Gating strategies for *in vivo* and *in vitro* infections and MHC II expression for infected BMDMs. A)** Gating strategy for Mtb-infected lung 14 dpi with msfYFP-H37Rv. **B)** Gating strategy for Mtb-infected mexAMs, and **C)** Gating strategy for Mtb-infected BMDMs with mCherry-H37Rv. **D)** MHC II MFI for bystander and Mtb-infected BMDMs 1 dpi, pre-treated with or without L-sulf for 24 hours. **D)** Representative of more than 3 independent experiments. **D)** Two-way ANOVA Turkey post-test. *p<0.05, **p<0.01, ***p<0.001, ****p<0.0001.

**Figure S2: NRF2 protein in AMs and BMDMs and baseline levels of MHC II genes in WT and NRF2^-/-^ mexAMs. A, B)** NRF2 protein 24 hours after LPS stimulation and/or L-sulf pre-treatment in WT AMs **(A)** and BMDMs (**B)**. **C, D)** *Ciita* **(C)** and *H2-Ab1* **(D)** expression in WT and NRF2^-/-^ mexAMs. **A, B)** Representative of 1 independent experiment or **C,D)** more than 4 independent experiments. **C,D)** Unpaired Student’s t-test. *p<0.05, **p<0.01, ***p<0.001, ****p<0.0001.

**Figure S3: *In vitro* gating strategies for AMs and BMDMs. A)** Gating strategy for AMs. Gating strategy for BMDMs. **C)** MHC II expression in untreated and LPS stimulated AMs. **D)** MHC II expression in untreated and LPS stimulated BMDMs.

**Figure S4: NRF2 inhibits MHC II expression in BMDMs, but not BMDCs, following PAMP stimulation in the presence of NRF2 agonist, L-sulf. A)** MHC II MFI in WT and NRF2^-/-^ BMDMs pre-treated with or without L-sulf for 24 hours, followed by PAMP stimulation (LPS or 2’3’-cGAMP) for an additional 24 hours. **B)** Percent inhibition of MHC II expression in NRF2^-/-^ BMDMs after L-sulf treatment compared to WT BMDMs across PAMP stimulations. **C)** MHC II MFI for BMDCs pre-treated with L-sulf prior to LPS stimulation. **A-C)** Representative of 2-4 independent experiments per condition. **A, C)** Two-way ANOVA with Tukey post-test. **B)** Unpaired Student’s t-test. *p<0.05, **p< 0.01, ***p< 0.001, ****p< 0.0001.

**Figure S5: Gating strategy for CD4^+^ T cell-mexAM co-culture after *in vitro* Mtb infection and BMDM co-culture. A)** Gating strategy for WT and NRF2^-/-^ mexAMs co-cultured with CD4^+^ T cells after Mtb infection. **B)** IFNγ production following 72 hour co-culture of C7 T cells and WT or NRF2^-/-^ BMDMs loaded with 0.1-10µM ESAT6_3-17_ peptide in the presence of L-sulf. B) Representative of 3 independent experiments. **B)** Two-way ANOVA with Tukey post-test. ****p< 0.0001.

**Figure S6: Summary graphic.** Expression of NRF2 in AMs limits MHC II expression. NRF2^-/-^ AMs have higher *Ciita* and *H2-Ab1* gene expression than WT AMs, leading to increased internal and surface MHC II protein. In co-culture with CD4^+^ T cells specific for Mtb antigens, Mtb-infected NRF2^-/-^ AMs promote higher T cell activation and production of IFNγ than WT AMs.

