## Supplementary material for "NRF2 inhibition of alveolar macrophage MHC II expression during *Mycobacterium tuberculosis* infection": Figures S1-6

**Supplementary Materials**

Figure S1: Gating strategies for *in vivo* and *in vitro* infections and MHC II expression for infected BMDMs.

Figure S2: NRF2 protein in AMs and BMDMs and baseline levels of MHC II genes in WT and NRF2^-/-^ mexAMs.

Figure S3: *In vitro* gating strategies for AMs and BMDMs

Figure S4: NRF2 inhibits MHC II expression in BMDMs, but not BMDCs, following PAMP stimulation in the presence of NRF2 agonist, L-sulf.

Figure S5: Gating strategy for CD4^+^ T cell-mexAM co-culture after *in vitro* Mtb infection and BMDM co-culture.

Figure S6: Summary graphic.

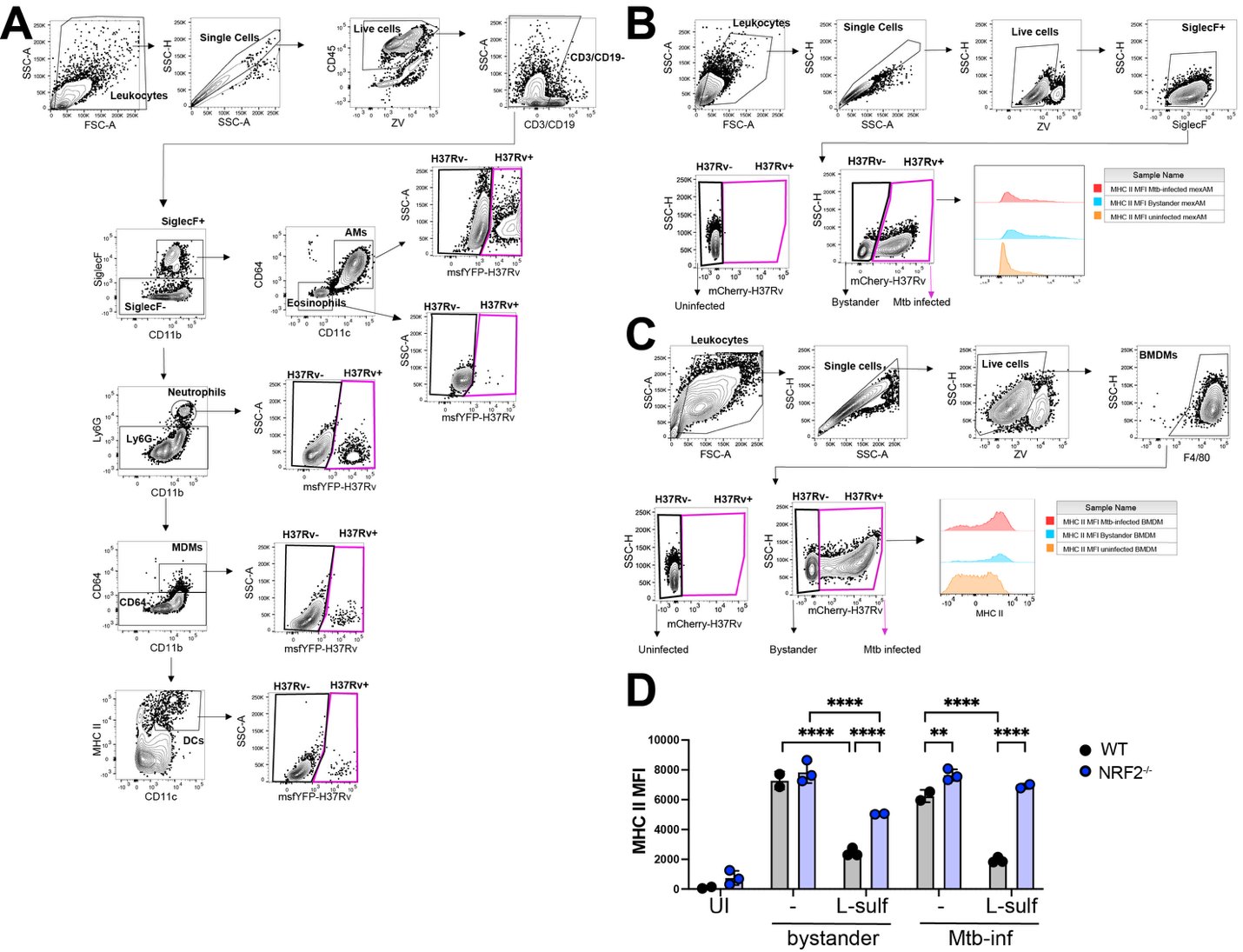

**Figure S1: Gating strategies for *in vivo* and *in vitro* infections and MHC II expression for infected BMDMs. A)** Gating strategy for Mtb-infected lung 14 dpi with msfYFP-H37Rv. **B)** Gating strategy for Mtb-infected mexAMs, and **C)** Gating strategy for Mtb-infected BMDMs with mCherry-H37Rv. **D)** MHC II MFI for bystander and Mtb-infected BMDMs 1 dpi, pre-treated with or without L-sulf for 24 hours. **D)** Representative of more than 3 independent experiments. **D)** Two-way ANOVA Turkey post-test. *p<0.05, **p<0.01, ***p<0.001, ****p<0.0001.

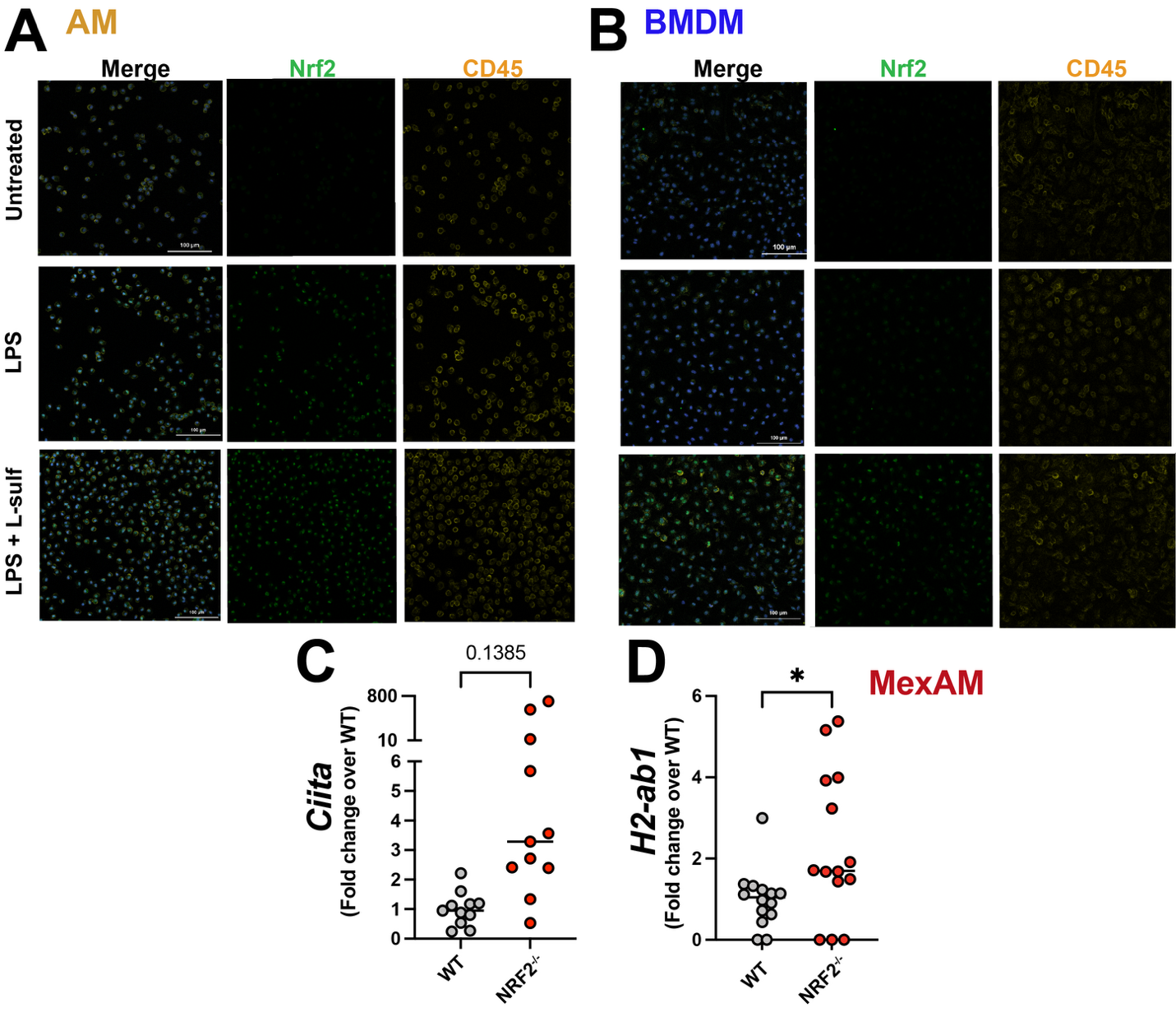

**Figure S2: NRF2 protein in AMs and BMDMs and baseline levels of MHC II genes in WT and NRF2^-/-^ mexAMs. A, B)** NRF2 protein 24 hours after LPS stimulation and/or L-sulf pre-treatment in WT AMs **(A)** and BMDMs (**B)**. **C, D)** *Ciita* **(C)** and *H2-Ab1* **(D)** expression in WT and NRF2^-/-^ mexAMs. **A, B)** Representative of 1 independent experiment or **C,D)** more than 4 independent experiments. **C,D)** Unpaired Student’s t-test. *p<0.05, **p<0.01, ***p<0.001, ****p<0.0001.

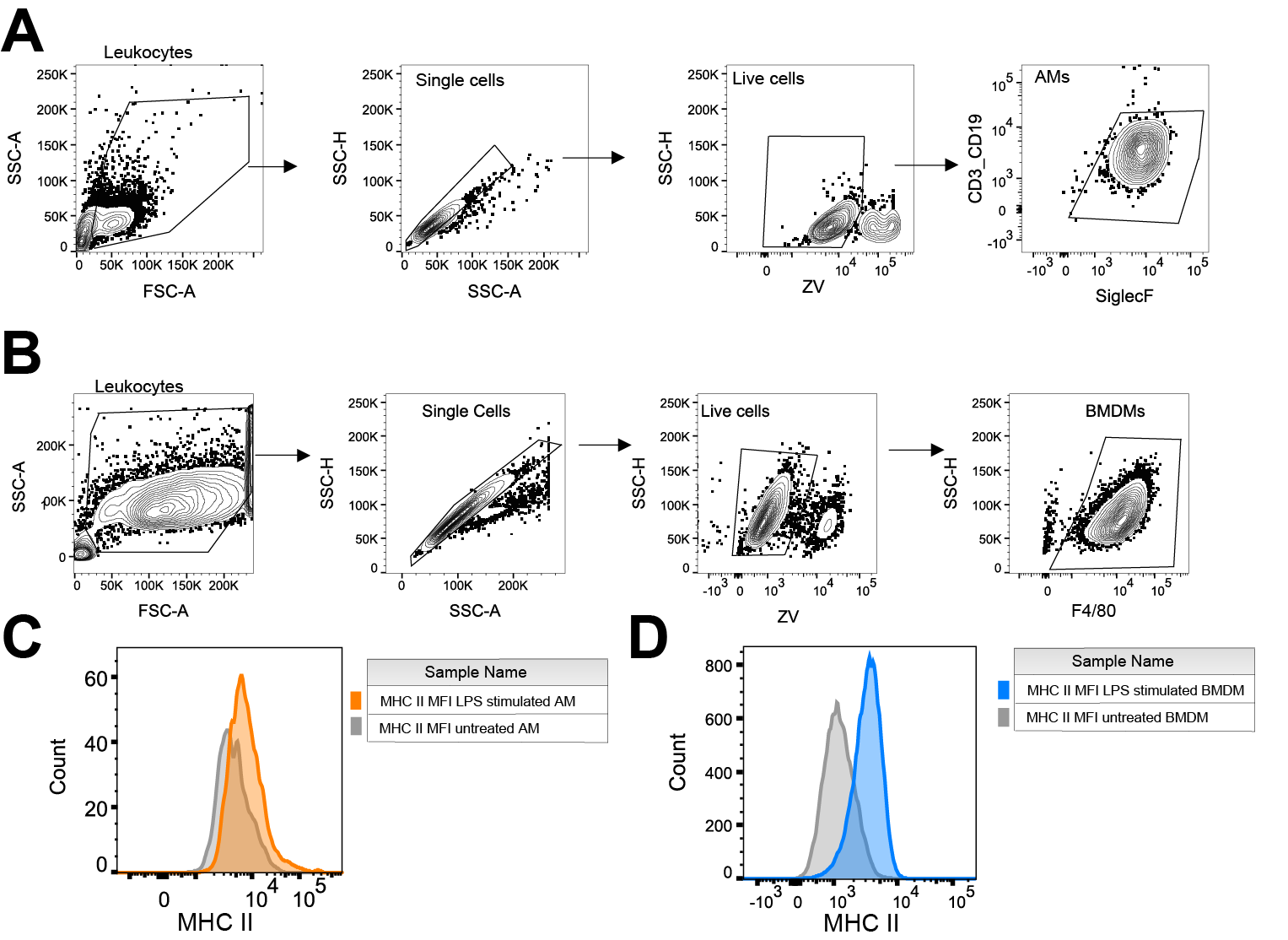

**Figure S3: *In vitro* gating strategies for AMs and BMDMs. A)** Gating strategy for AMs. **B)** Gating strategy for BMDMs. **C)** MHC II expression in untreated and LPS stimulated AMs. **D)** MHC II expression in untreated and LPS stimulated BMDMs.

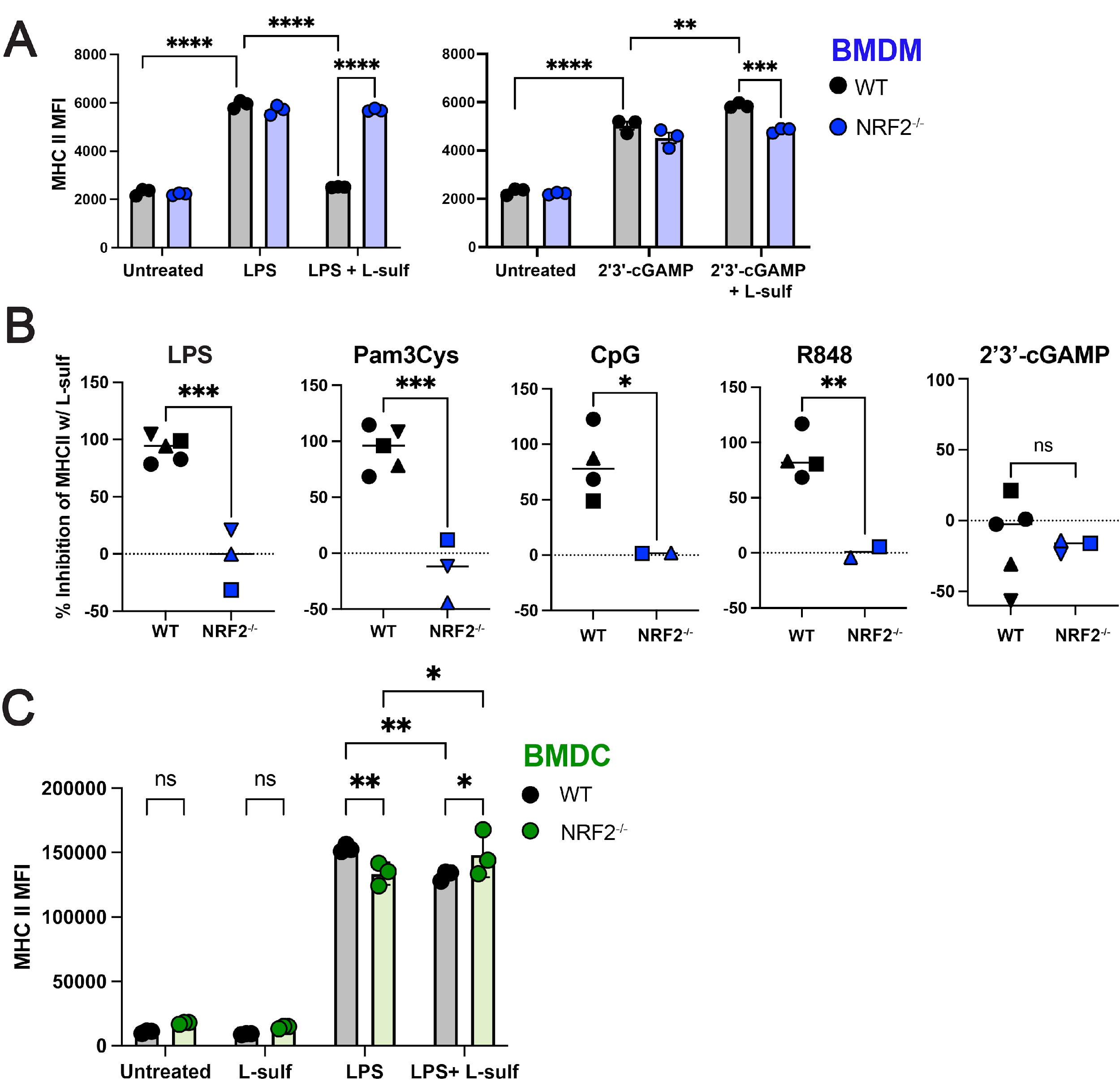

**Figure S4: NRF2 inhibits MHC II expression in BMDMs, but not BMDCs, following PAMP stimulation in the presence of NRF2 agonist, L-sulf. A)** MHC II MFI in WT and NRF2^-/-^ BMDMs pre-treated with or without L-sulf for 24 hours, followed by PAMP stimulation (LPS or 2’3’-cGAMP) for an additional 24 hours. **B)** Percent inhibition of MHC II expression in NRF2^-/-^ BMDMs after L-sulf treatment compared to WT BMDMs across PAMP stimulations. **C)** MHC II MFI for BMDCs pre-treated with L-sulf prior to LPS stimulation. **A-C)** Representative of 2-4 independent experiments per condition. **A, C)** Two-way ANOVA with Tukey post-test. **B)** Unpaired Student’s t-test. *p<0.05, **p< 0.01, ***p< 0.001, ****p< 0.0001.

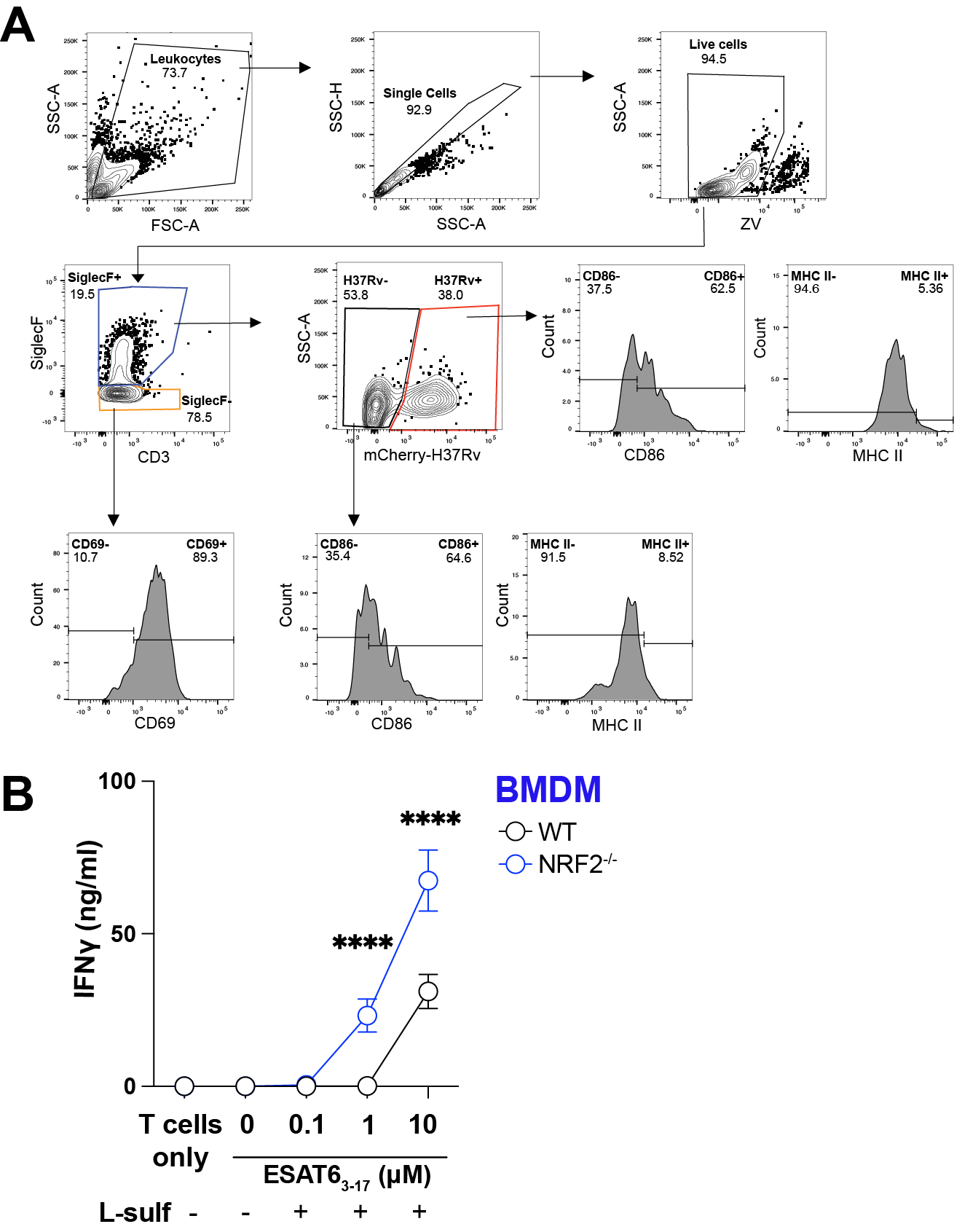

**Figure S5: Gating strategy for CD4^+^ T cell-mexAM co-culture after *in vitro* Mtb infection and BMDM co-culture. A)** Gating strategy for WT and NRF2^-/-^ mexAMs co-cultured with CD4^+^ T cells after Mtb infection. **B)** IFNγ production following 72 hour co-culture of C7 T cells and WT or NRF2^-/-^ BMDMs loaded with 0.1-10uM ESAT6_3-17_ peptide in the presence of L-sulf. B) Representative of 3 independent experiments. **B)** Two-way ANOVA with Tukey post-test. ****p< 0.0001.

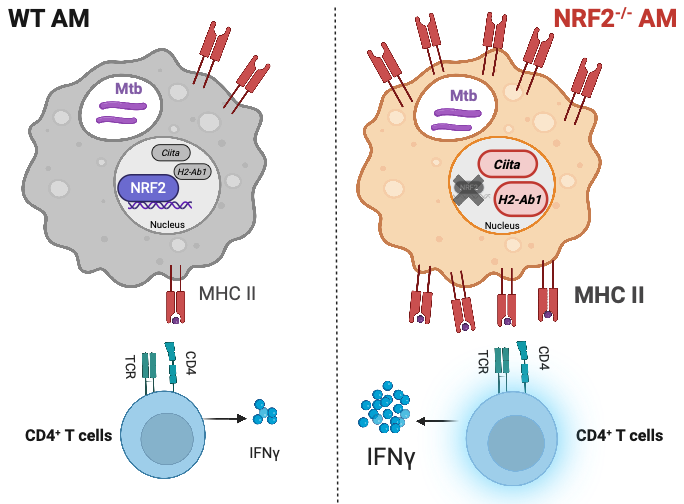

**Figure S6: Summary graphic.** Expression of NRF2 in AMs limits MHC II expression. NRF2^-/-^ AMs have higher *Ciita* and *H2-Ab1* gene expression than WT AMs, leading to increased internal and surface MHC II protein. In co-culture with CD4^+^ T cells specific for Mtb antigens, Mtb-infected NRF2^-/-^ AMs promote higher T cell activation and production of IFNγ than WT AMs.
